# Large excess capacity of glycolytic enzymes in *Saccharomyces cerevisiae* under glucose-limited conditions

**DOI:** 10.1101/2022.07.06.498936

**Authors:** Pranas Grigaitis, Bas Teusink

## Abstract

In Nature, microbes live in very nutrient-dynamic environments. Rapid scavenging and consumption of newly introduced nutrients therefore offer a way to outcompete competitors. This may explain the observation that many microorganisms, including the budding yeast *Saccharomyces cerevisiae*, appear to keep “excess” glycolytic proteins at low growth rates, i.e. the maximal capacity of glycolytic enzymes (largely) exceeds the actual flux through the enzymes. However, such a strategy requires investment into preparatory protein expression that may come at the cost of current fitness. Moreover, at low nutrient levels, enzymes cannot operate at high saturation, and overcapacity is poorly defined without taking enzyme kinetics into account.

Here we use computational modeling to suggest that in yeast the overcapacity of the glycolytic enzymes at low specific growth rates is a genuine excess, rather than the optimal enzyme demand dictated by enzyme kinetics. We found that the observed expression of the glycolytic enzymes did match the predicted optimal expression when *S. cerevisiae* exhibits mixed respiro-fermentative growth, while the expression of tricarboxylic acid cycle enzymes always follows the demand. Moreover, we compared the predicted metabolite concentrations with the experimental measurements and found the best agreement in glucose-excess conditions. We argue that the excess capacity of glycolytic proteins in glucose-scarce conditions is an adaptation of *S. cerevisiae* to fluctuations of nutrient availability in the environment.

## Introduction

It is a common observation that microorganisms, growing in carbon-limited conditions, possess reserve glycolytic capacity, which means that the abundances, and, subsequently, the maximal reaction rates of glycolytic enzymes largely exceed *in vivo* fluxes. Such a phenomenon was observed in various microbes, such as *Escherichia coli* (Schmidt *et al*, 2016), *Bacillus subtilis* (Buffing *et al*, 2018), and *Lactococcus lactis* (Goel *et al*, 2015). Among them, the budding yeast *Saccharomyces cerevisiae* is renowned for its ability to consume glucose at high rates (Blank & Sauer, 2004).

Although the fluxes through glycolytic enzymes are very low at strong glucose limitation in S. *cerevisiae*, various studies have reported that enzyme activities (Van Hoek *et al*, 1998) and mRNA levels (Sierkstra *et al*, 1992) barely change for most glycolytic enzymes across a broad span of dilution rates in glucose-limited chemostats. Additionally, we recently reported a stable fraction of total S. *cerevisiae* proteome occupied by glycolytic enzymes in glucose-limited chemostats spanning *D* = 0.20 to 0.34 *h*^-1^ ((Elsemman *et al*, 2022), Figure 1a, 1b). Combined, these are indeed remarkable observations, considered that in some of the aforementioned studies, the specific glucose uptake rate varied up to ca. 15-fold across conditions. Overall, there seems to be a profound reserve capacity of glycolytic enzymes in *S. cerevisiae* growth in glucose-scarce environments.

**Figure 1.**
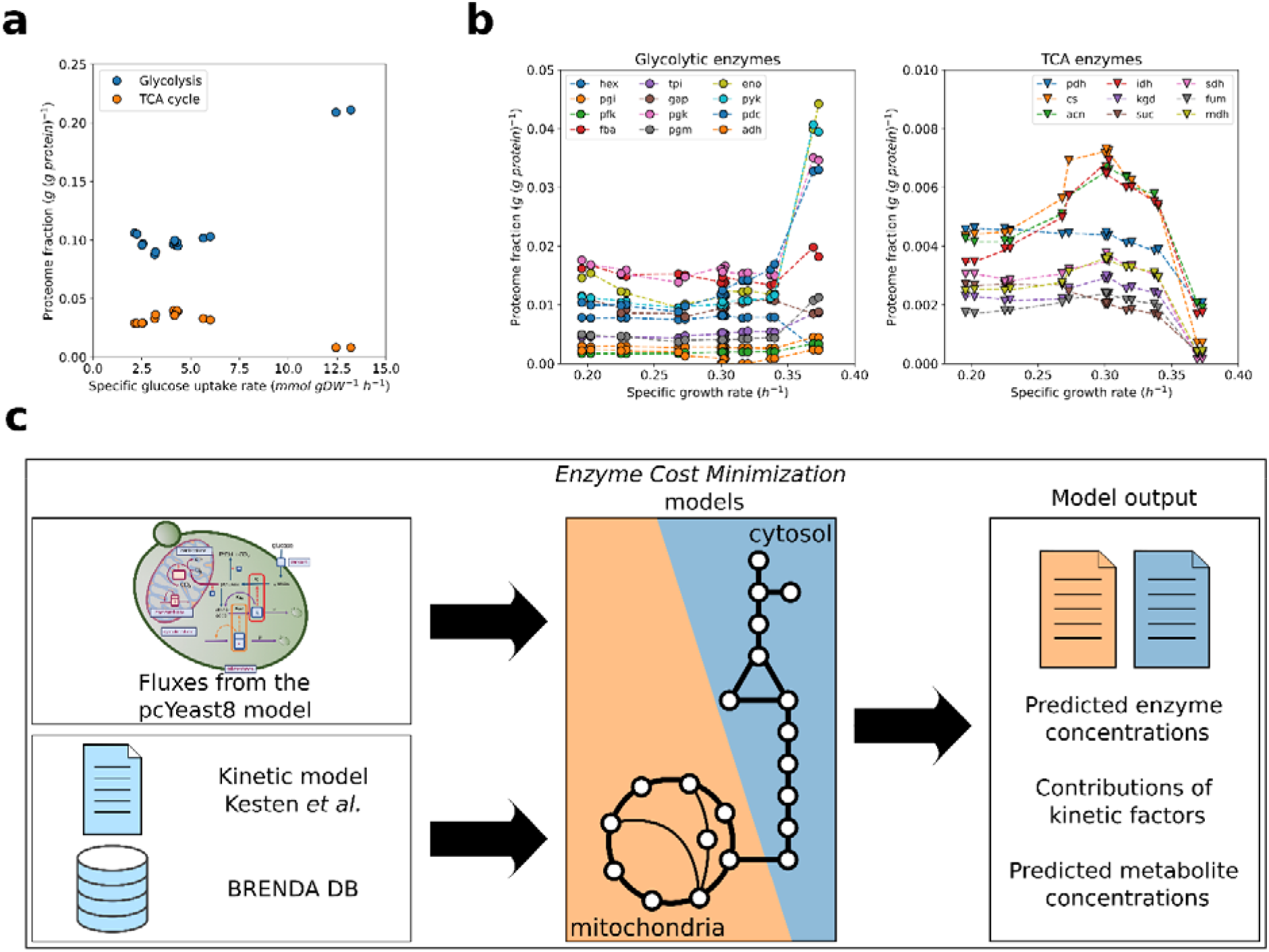
Observed expression levels of enzymes of central carbon metabolism in *S. cerevisiae* and our computational approach. **a**. Proteome mass fractions of glycolytic and TCA cycle enzymes as a function of specific glucose uptake rate. **b.** The breakdown of the proteome mass fractions for different glycolytic (left) and TCA cycle (right) enzymes (isozymes lumped together) as a function of specific growth rate. **c.** The modeling approach used in this study. We computed the steady-state fluxes for the glucose-limited and glucose-excess conditions described in (Elsemman *et al*, 2022) in the *pcYeast8* proteome-constrained model of *S. cerevisiae* metabolism (Grigaitis *et al*., unpublished) and used that as an input for the *Enzyme Cost Minimization* models (see Methods for details of model construction and Figure S1 for the distribution of kinetic values). Quantitative proteomics data for (**a-b**) from (Elsemman *et al*, 2022).

The (over)abundance of glycolytic proteins may be rationalized as a short-term adaptation to (non)periodic oscillations of glucose levels, e.g. sudden availability (glucose pulse) or feast/famine cycles, which are fairly frequent in natural environments. The increased consumption of glucose straight after a glucose pulse was shown to be protein synthesis-independent (Reijenga *et al*, 2005), and is driven by metabolic regulation, reflected in increased concentrations of glycolytic intermediates (Wu *et al*, 2006; van den Brink *et al*, 2008; Suarez-Mendez *et al*, 2017). Only tens of minutes post-pulse, increased mRNA levels of glycolytic enzymes (van den Brink *et al*, 2008; Dikicioglu *et al*, 2011) and synthesis of low-affinity glucose transporters (Buziol *et al*, 2008) are observed.

However, whether the glycolytic capacity is really in excess, is not clear to the day. Low glycolytic flux in these cases is caused by low extracellular glucose, and hence low pathway substrate. Consequently, there is a low thermodynamic driving force and hence metabolite concentrations for all subsequent enzymes. So perhaps the high expression of enzymes reflects *the* optimal enzyme levels needed to accommodate the low fluxes, given enzyme reversibility and undersaturation.

To answer this question, here we use metabolic models of *S. cerevisiae* to quantify the enzyme demand needed to run central carbon metabolism at different glucose levels corresponding fluxes (Figure 1c). We used the *Enzyme Cost Minimization* (ECM) method to compute enzyme demands and compared them with quantitative proteomics data. We find that the computed enzyme demands match the experimental measurements in the high flux, respiro-fermentative regime, while at the low flux, fully-respiratory growth, kinetic effects do not fully explain the observed expression levels. We thus conclude that the *S. cerevisiae* cells indeed do have an excess capacity of glycolytic enzymes, which may provide advantage for the short-term adaptation in dynamic environments.

## Results

### Overview of the modeling approach

The *Enzyme Cost Minimization* (ECM) method computes the minimal enzyme demand required to support given fluxes in metabolic networks, based on the kinetic properties of the enzymes and thermodynamic properties of the reactions (Gibbs free energy change) (Noor *et al*, 2016). The method takes steady-state flux values as input. Then, the weighted sum of enzyme demands is minimized, with the enzyme concentrations and natural logarithms of metabolite concentrations as decision variables. The overall enzyme demand is defined as a combination of the minimal demand, i.e. the ratio between the flux through the enzyme and its turnover value (*k_cat_*), and two aspects of enzyme kinetics: reaction reversibility (thermodynamic driving force), and enzyme saturation. We adapted the ECM model of *Escherichia coli* central carbon metabolism (CCM) to reconstruct two pathway models of *S. cerevisiae*: the cytosolic CCM (glycolysis and pentose phosphate pathway), and mitochondrial CCM (tricarboxylic acid [TCA] cycle) (Methods). In this study, we considered the 7 conditions of growth on glucose reported in (Elsemman *et al*, 2022): 6 glucose-limited chemostats (0.20, 0.23, 0.27, 0.30, 0.32, 0.34 *h*^-1^), and a culture in excess of glucose (*μ* = 0.37 to 0.40 *h*^-1^). We then simulated the models to predict the enzyme levels and metabolite concentrations (Figure 1c).

### Glycolytic proteins are expressed in excess in respiratory growth

We first asked what the predicted enzyme demands are at different growth rates and compared them to the experimental data (Figure 2a, 2b). We first considered the “minimal” enzyme demand (thus assuming operation of enzymes at maximal capacity) and observed a large discrepancy between the measured enzyme levels vs. the minimal demand, even at glucose-excess conditions. Inclusion of the effect of enzyme kinetics resulted in a much higher demand for enzyme, as expected. The key finding is that at fully respiratory growth (*D* < *D_crit_* = 0.28 *h*^-1^), the measured abundance of pooled glycolytic enzymes could not be fully explained by the combination of the kinetic effects (Figure 2a). Meanwhile, the predicted demand seemed to correspond well with the experimental measurements when respiro-fermentative growth is observed. On the contrary, the predicted demand of the total of TCA cycle enzymes (Figure 2b) matched the experimental measurements well at all growth rates.

**Figure 2.**
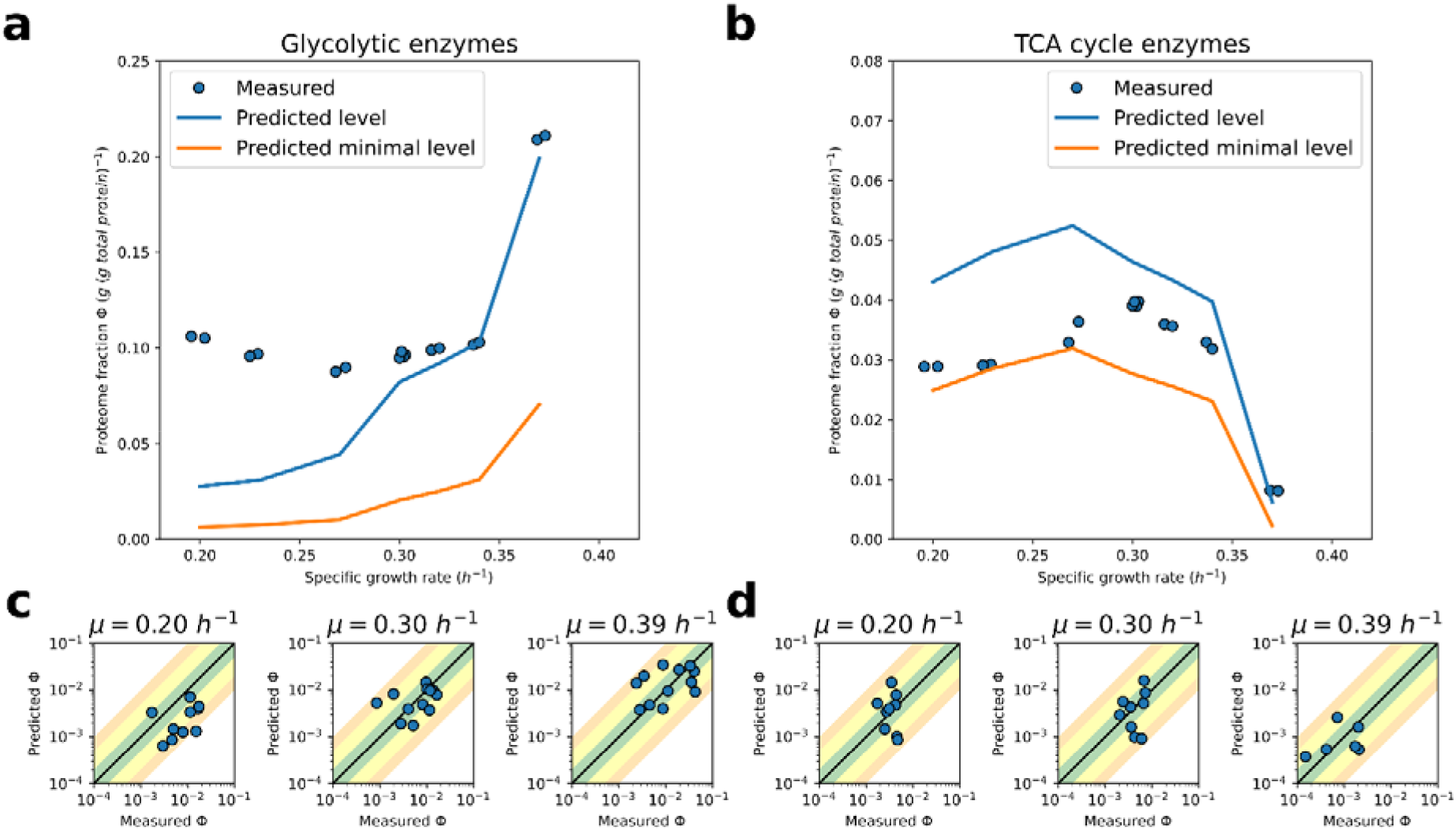
Predicted proteome mass fractions of the glycolytic and TCA cycle enzymes. **a-b.** Predicted enzyme demands (minimal demand,, orange line; demand including kinetic aspects, blue line) of glycolytic (a) and TCA cycle enzymes (b) as a function of the specific growth rate. **c-d.** Predicted (including kinetic aspects) proteome fractions of individual glycolytic (c) and TCA cycle (d) enzymes at 3 different specific growth rates. The green, yellow, and orange shaded regions represent the agreement of predicted vs. measured fractions in the 2-, 5-, and 10-fold range, respectively. Quantitative proteomics data from (Elsemman *et al*, 2022).

We then looked at the demands at the individual enzyme level (Figure 2c, 2d), where we observed more discrepancies. As the fluxes increased (=increasing growth rate), we observed that the predicted demand of glycolytic enzymes generally increased (Figure 2c), but their demand was almost universally underpredicted by a large margin at lower growth rates. The latter conclusion is even more evident comparing the minimal predicted vs. observed enzyme abundance (Figure S2), where we observed the underprediction of minimal levels to be generally in the range of 5- to 50-fold for many of the glycolytic enzymes. As expected, the major contribution to enzyme demands comes from the undersaturation of glycolytic enzymes (Figure S3), with a substantial contributions of enzyme reversibility for the isomerases (glucose 6-phosphate isomerase, and triose phosphate isomerase), as well as fructose 1,6-bisphosphate aldolase.

It should be noted here that the predicted enzyme demands are particularly sensitive to the *k_cat_* values: the enzyme demand, accounted for kinetic effects, are multipliers of the minimal enzyme demand, rather than additive increments. The predicted demands of individual TCA cycle enzymes (Figure 2d) seem to be influenced by the uncertainty of the *k_cat_* values. Yet, unlike the glycolytic enzymes, there was no systemic underprediction of the enzyme demands for TCA enzymes. This thus suggests that the “combined” prediction of the proteome fractions (Figure 2b) is a robust outcome, although for individual enzymes we observed under- and overpredicted demands. Moreover, a distinct property of the predicted TCA cycle enzyme demands is that their minimal predicted demand was already a good approximation of their expression levels (Figure 2b, Figure S4), with a substantially smaller contribution to the overall demand coming from enzyme (under)saturation (Figure S4) due to generally lower *K_M_* values (Figure S1). With this, we conclude that the demand of the TCA cycle enzymes seems to be set by the flux through these enzymes.

### Prediction of metabolite levels suggests optimal growth at glucose batch conditions

The key feature of the *Enzyme Cost Minimization* method, compared to other optimization-based metabolic modeling approaches, is that the natural logarithms of the concentrations of metabolites are optimization variables as well. Thus, alongside enzyme demands, we can describe the predicted metabolic states in terms of the steady-state concentrations of metabolites (Figure 3).

**Figure 3.**
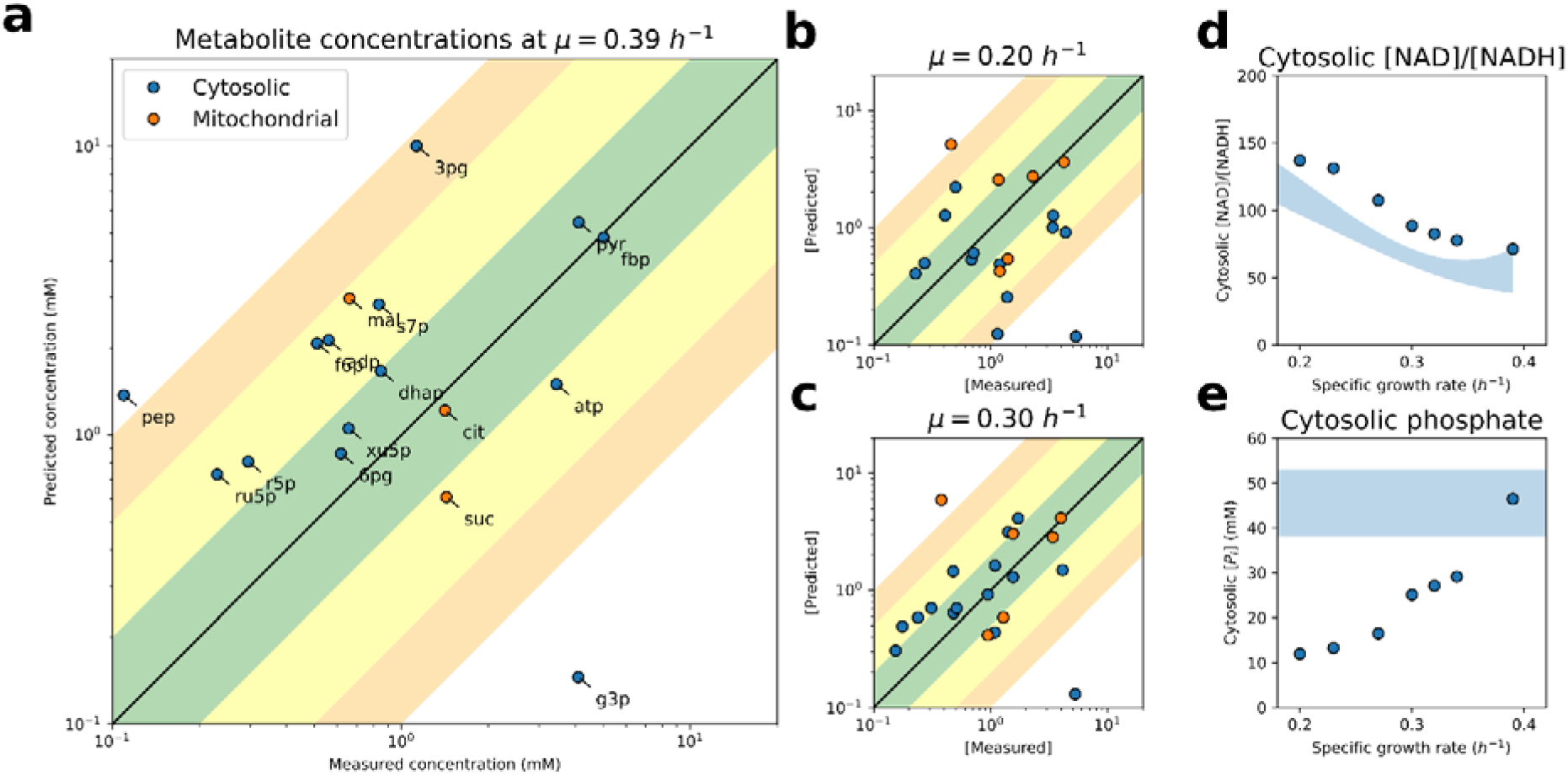
Comparison of predicted metabolite levels at different specific growth rates. **a-c.** Predicted metabolite concentrations at the glucose-excess (a) and glucose-limited (b-c) conditions. The green, yellow, and orange shaded regions represent the agreement of predicted vs. measured levels in the 2-, 5-, and 10-fold range, respectively. **d-e.** Predicted cytosolic [NAD]/[NADH] ratio (d) and cytosolic phosphate concentration (e). Shaded regions in (d-e) represent the 3^rd^ degree polynomial fit, or experimentally determined range of the experimental measurements, in (d) and (e), respectively. Metabolite concentration data from (Canelas *et al*, 2011).

The predicted metabolite concentrations (Figure 3a–3c) were, as for the enzyme demands, in best alignment with experimental data (Canelas *et al*, 2011) for the glucose-limited respiro-fermentative (Figure 3b) and glucose-excess conditions (Figure 3a). We also compared the predicted cytosolic [NAD]/[NADH] ratio, and found a good qualitative agreement with experimental data at all growth rates (Figure 3d). Eventually we observed that the predicted cytosolic [NAD]/[NADH] ratio and the concentration of cytosolic phosphate were in a good quantitative agreement with experimental data for the batch conditions (Figure 3d-e). Based on this, we suggest that at glucose-excess conditions, most of the enzymes in yeast CCM are expressed and operate optimally.

## Discussion

In this study, we sought to elucidate the nature of the perceived overcapacity of glycolytic enzymes, which is a trait observed in many microorganisms. Here, we focused on *S. cerevisiae* for our analysis, and compared enzyme demands, predicted by computational models, with experimental measurements. We found that, at low growth rates, the kinetic factors of enzyme catalysis (enzyme reversibility and undersaturation) explain only a fraction of the observed protein pool, pointing at a genuine excess capacity of enzymes, i.e. above the optimal level needed to support the steady-state flux under that condition (Figure 2a). One could reason that the allocation of additional resources to glycolysis is a preparatory (dynamic) resource allocation strategy in glucose-scarce environments, as was also suggested for ribosomal proteins (Metzl-Raz *et al*, 2017).

Under glucose-limited conditions, glucose transport seems to be *the* constraint which actively limits growth (Elsemman *et al*, 2022) as the response of prolonged cultivation in glucose-limited conditions is the duplication of glucose transporter loci (Brown *et al*, 1998), only later to be accompanied by a decrease of fermentative capacity (Jansen *et al*, 2005). Thus, at dynamically changing conditions, the selection pressure is rather on scavenging the limiting nutrients, and suboptimal (at that condition) allocation of resources is unlikely to be outcompeted by individuals whose expression of glycolytic enzymes matches the demand set by the glycolytic flux. This hypothesis is supported by computational modeling, as we observed that increased expression of gratuitous protein does not affect the predicted growth rate in nutrient-limited conditions ((Elsemman *et al*, 2022), Grigaitis *et al*., unpublished). It was shown that perturbations to the proteome composition have no influence on the physiology of glucose-limited chemostat cultures, e.g. when mitochondrial biosynthesis was increased by overexpression of transcription factor Hap4 (Maris *et al*, 2001).

Yet the situation changes in respiro-fermentative growth and/or glucose-excess conditions (Figure 2a, 2c). There, the predicted enzyme demand matches experimentally determined expression, and the ca. 2-fold increase in glycolytic flux between the last chemostat point (*μ* = 0.34 *h*^-1^) and the glucose excess (*μ* = 0.39 *h*^-1^) is followed by a similar increase in the proteome fraction, allocated to the glycolytic enzymes – predicted and measured alike. Previously we proposed that the growth in glucose-excess conditions is proteome-limited (Elsemman *et al*, 2022) and thus defined by optimal proteome allocation: expression of gratuitous proteins (not contributing to growth) results in a decrease of the maximal growth rate (Kafri *et al*, 2016). The quantitative agreement of the predicted enzyme demands (Figure 2a) and metabolite concentrations (Figure 3a, 3e) in glucose-excess conditions further supports this conclusion.

Unlike the glycolytic enzymes, we observed that the demand of the TCA cycle enzymes, in most cases, followed the demand set by flux (Figure 2b, 2d). Our findings support the previous observations, such as the idea that metabolically-active organelles (in this case, cytosol vs. mitochondria) compete for resources in yeast. As an example, a major rearrangement of cell composition happens during the glucose-ethanol diauxic shift: (Di Bartolomeo *et al*, 2020) reported a ca. 7-fold increase in the cell volume occupied by mitochondria, between the glucose- (pre-shift) and ethanol-excess (post-shift) conditions. As ethanol enters the cell metabolism primarily through the TCA cycle, and growth on ethanol requires high fluxes through the TCA cycle to assimilate enough ethanol, such a transition points to a demand-driven allocation of resources to mitochondria.

To conclude, we argue that the excess capacity of glycolytic proteins in glucose-scarce conditions is an adaptation to fluctuations of nutrient availability in the environment. Being able to swiftly consume as much glucose as possible provides an advantage over competitors and happens without new protein synthesis, as an increase in uptake of glucose happens in the time-scale of seconds and minutes post-pulse (van den Brink *et al*, 2008). We here considered only one microbe as an example, *Saccharomyces cerevisiae*, yet expect this excess capacity of enzymes to be a unifying trait of nutrient-limited growth of microbes.

## Methods

### Data Collection

Label-free quantitative proteomics data from yeast cultures, grown in glucose-limited chemostats at 6 different dilution rates, as well as batch cultures with excess glucose, were acquired from (Elsemman *et al*, 2022). Kinetic data (*k_cat_* and *K_M_* values) for the wild-type enzymes of *S. cerevisiae* at conditions closest to the physiological ones (temperature 30 °C, pH 7 etc.) were gathered from BRENDA database (Chang *et al*, 2021) (Figure S1). Information on protein masses, stoichiometry of complexes were collected from UniProt (The UniProt Consortium *et al*, 2021).

### *Enzyme Cost Minimization* Model

*Enzyme Cost Minimization* (ECM) modeling framework (Noor *et al*, 2016) was used. The models of yeast central carbon metabolism (CCM) were reconstructed using of the model of *E. coli* CCM as a template, provided in the method paper (Noor *et al*, 2016). The model was curated as follows: reactions *ppc* and fbp from the template model were removed, and we added the ethanol fermentation branch (reactions *pdc* and *adh*), as well as the external- and internal NADH oxidases (*nde* and *ndi*, cytosolic and mitochondrial, respectively). To allow metabolites to acquire different concentration values in different compartments, the reconstructed CCM model was split into a cytosolic (glycolysis, pentose phosphate pathway, ethanol fermentation) and mitochondrial (TCA cycle) model. When there was no experimentally determined *K_M_* value, a default *K_M_* = 0.1 *mM* was assumed. The parameters of pH, pMg, and ionic strength were retained as in the *E. coli* model (7, 3, and 250 mM, respectively).

Concentrations of intracellular glucose and ethanol were fixed as follows: for glucose, the intracellular [glucose] was estimated from the data of glucose transport capacity at different dilution rates in glucose-limited chemostats/glucose batch cultures from (Diderich *et al*, 1999) using the computation provided in (Teusink *et al*, 1998) (Table S1). For ethanol, the intracellular concentration of 0.001 mM was assumed. Concentrations of all other metabolites were allowed to vary.

For every condition (as described in the Results, “Overview of the modeling approach”), a set of cytosolic and mitochondrial models was generated. Intracellular steady-state fluxes were computed by the *pcYeast8* model ((Elsemman *et al*, 2022), Grigaitis et al., unpublished) and used as input to the ECM models. Reactions, carrying zero flux in the *pcYeast8* model were removed altogether from the ECM models. The predicted enzyme concentrations in mM were converted to proteome fractions by assuming the fixed relationship of *V_cell_* = 1.7 *mL gDW*^-1^ and bulk protein composition in biomass: *f_p_* = 0.42436 *μ* + 0.35364 (*g protein* (*g biomass*)^-1^).

## Data Availability

Models, data & code to reproduce the results are provided on Zenodo [10.5281/zenodo.6801406] (Grigaitis, Pranas & Teusink, Bas, 2022).

## Acknowledgements

We thank Ursula Kummer for discussions and feedback on the results, and Elad Noor and Wolfram Liebermeister for the discussions regarding the Enzyme Cost Minimization method. The authors acknowledge the funding by Marie Skłodowska-Curie Actions ITN “SynCrop” (grant agreement No 764591) and NWO (NWO ERA-IB-2, project No 053.80.722).

## CRediT Author Role Statement

Pranas Grigaitis – data curation, formal analysis, investigation, methodology, resources, software, validation, visualization, writing – original draft, writing – review & editing; Bas Teusink – conceptualization, formal analysis, funding acquisition, project administration, supervision, validation, writing – review & editing.

