## Supplementary Material for "Large excess capacity of glycolytic enzymes in *Saccharomyces cerevisiae* under glucose-limited conditions"

**This document contains:**

Supplementary Figures 1-4

Supplementary References

**Separate supplementary files:**

Supplementary Table 1 (.xlsx)

### Supplementary Figures

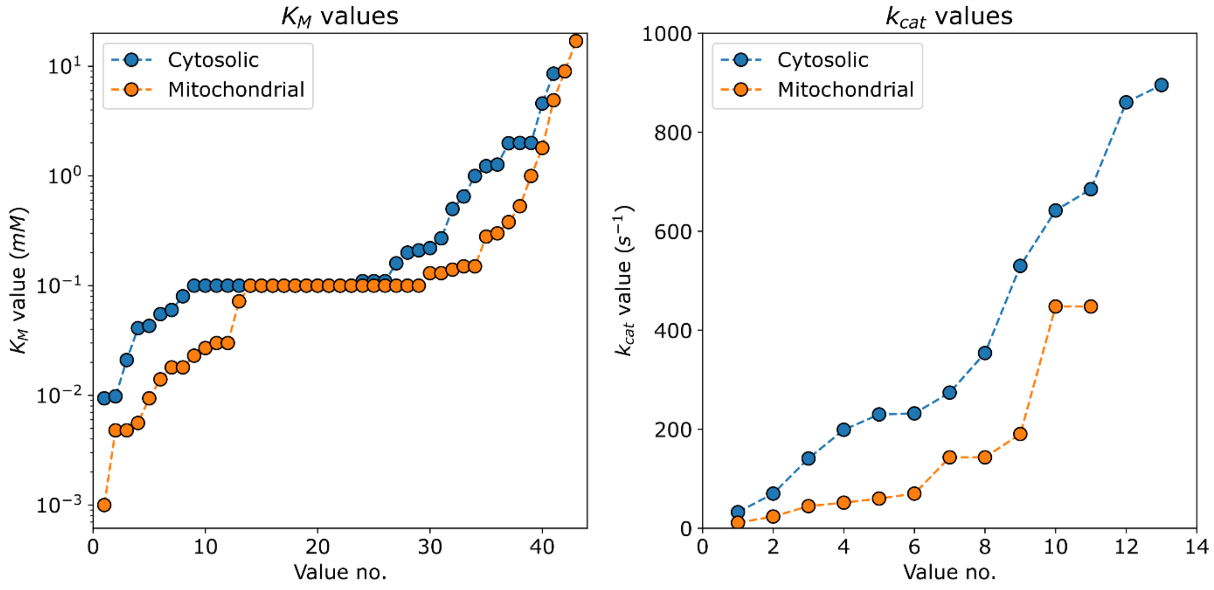


**Figure S1.** Comparison of the kinetic values in the cytosolic and mitochondrial *Enzyme Cost Minimization* models. The values were collected as described in the Methods section of the Main Text, ranked from lowest to highest, and plotted for both cytosolic (glycolysis) and mitochondrial (TCA cycle) enzymes. The plateau of $K_M$ values corresponds to the default value of $K_M = 0.1\ mM$.

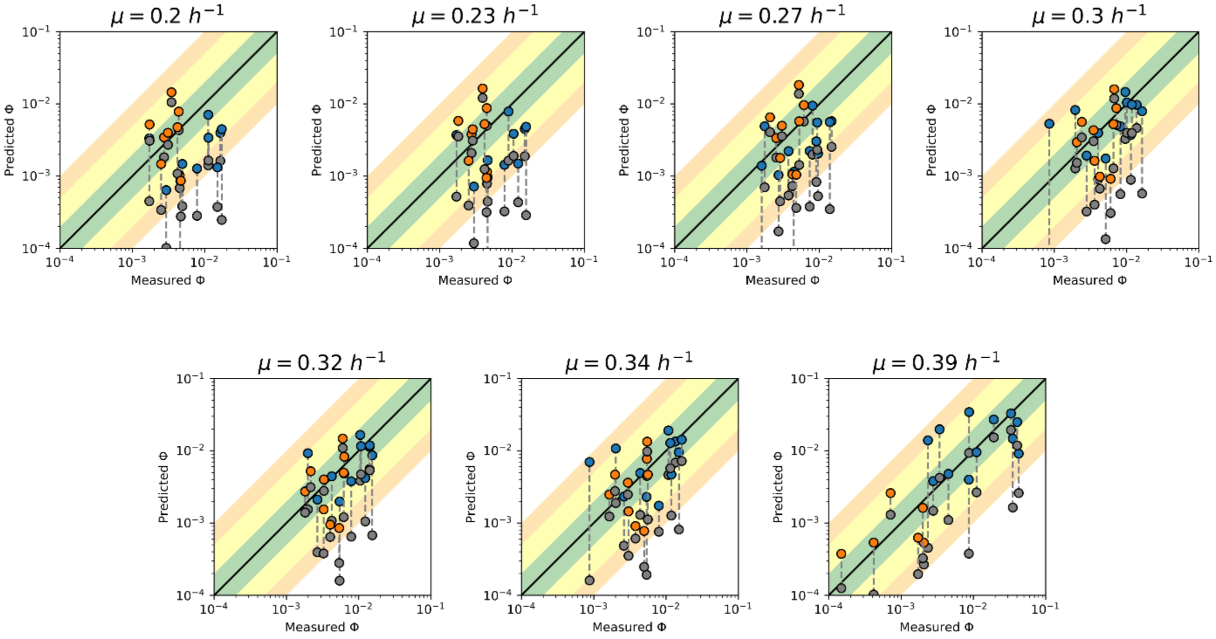


**Figure S2.** Predicted proteome fractions of individual glycolytic (blue points) and TCA cycle (orange points) enzymes at different specific growth rates. The grey points indicate the minimal predicted proteome fraction. The green, yellow, and orange shaded regions represent the agreement of predicted vs. measured fractions in the 2-, 5-, and 10-fold range, respectively. Quantitative proteomics data from (Elsemman et al., 2022).

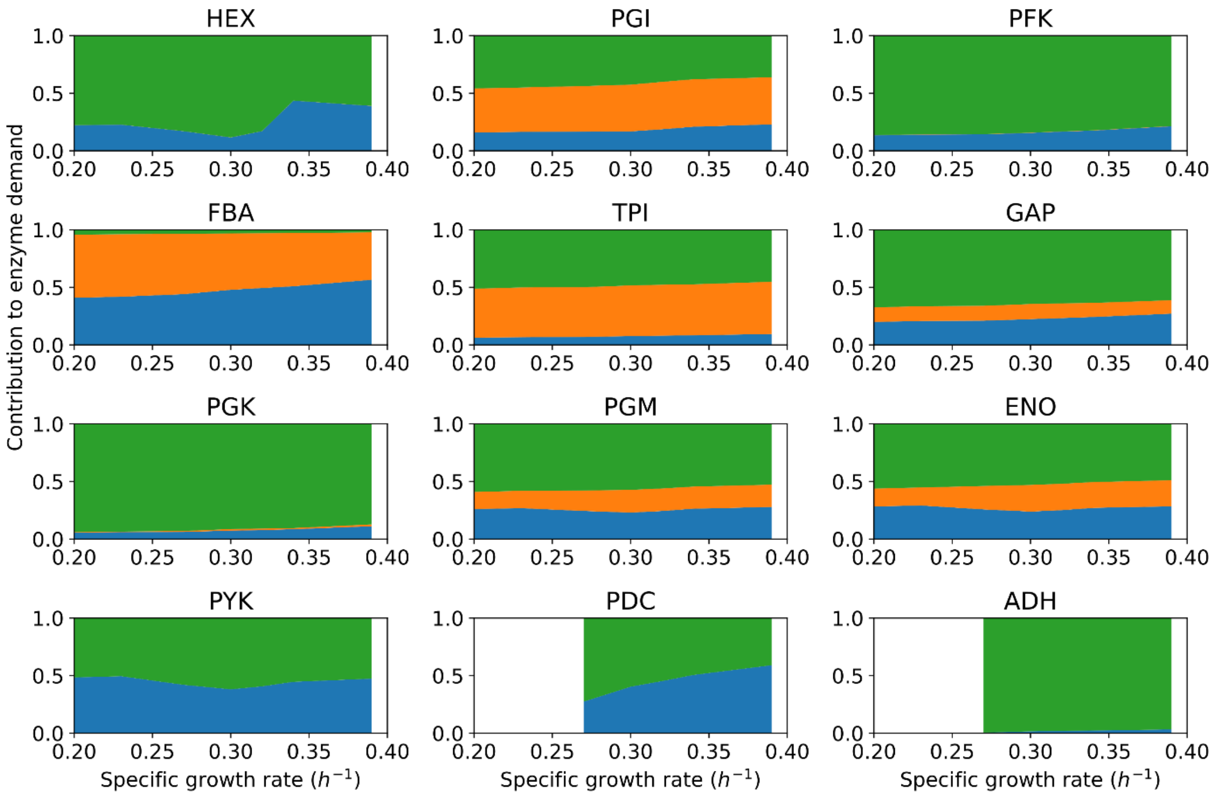


**Figure S3.** The contribution of different kinetic factors to the computed glycolytic enzyme demand (Figure 2) at different specific growth rates. Blue, demand by flux; orange, reversibility of enzymes; green, saturation of enzymes. Some of the trajectories are incomplete as these enzymes do not carry flux at some growth rates.

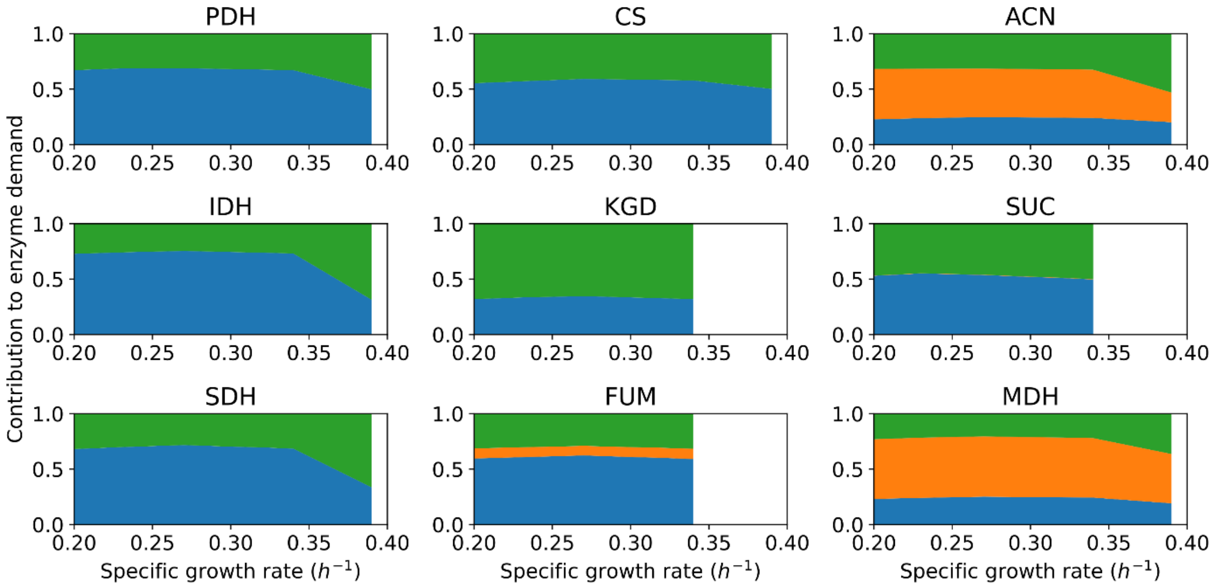


**Figure S4.** The contribution of different kinetic factors to the computed TCA cycle enzyme demand (Figure 2) at different specific growth rates. Blue, demand by flux; orange, reversibility of enzymes; green, saturation of enzymes. Some of the trajectories are incomplete as these enzymes do not carry flux at some growth rates.
